# Neglecting model selection alters phylogenetic inference

**DOI:** 10.1101/849018

**Authors:** Michael Gerth

## Abstract

Molecular phylogenetics is a standard tool in modern biology that informs the evolutionary history of genes, organisms, and traits, and as such is important in a wide range of disciplines from medicine to palaeontology. Maximum likelihood phylogenetic reconstruction involves assumptions about the evolutionary processes that underlie the dataset to be analysed. These assumptions must be specified in forms of an evolutionary model, and a number of criteria may be used to identify the best-fitting from a plethora of available models of DNA evolution. Using many empirical and simulated nucleotide sequence alignments, Abadi et al.^1^ have recently found that phylogenetic inferences using best models identified by six different model selection criteria are, on average, very similar to each other. They further claimed that using the model GTR+I+G4 without prior model-fitting results in similarly accurate phylogenetic estimates, and consequently that skipping model selection entirely has no negative impact on many phylogenetic applications. Focussing on this claim, I here revisit and re-analyse some of the data put forward by Abadi et al. I argue that while the presented analyses are sound, the results are misrepresented and in fact - in line with previous work - demonstrate that model selection consistently leads to different phylogenetic estimates compared with using fixed models.

## MAIN TEXT

To assess the impact of different model selection criteria on phylogenetic accuracy, Abadi et al. acquired 7,200 nucleotide alignments from various databases (empirical dataset), from which three equal-sized datasets with increasing complexity were simulated under common nucleotide substitution models (datasets c_0_–c_2_). A smaller dataset was simulated under a codon substitution model (c_3_). For all alignments across datasets, maximum likelihood estimations were performed using the “best” models determined by six different selection criteria, and the fixed models GTR+I+G4 and JC. Differences in topologies were recorded using Robinson-Foulds distances or by simply counting non-identical trees. Abadi et al.’s claim that model selection is redundant stems mainly from three observations: 1) Trees inferred under different model selection criteria are often identical; 2) The proportion of correctly inferred topologies is highly similar between all model selection criteria and fixed models; 3) Topological distances between trees inferred under any strategy are also very similar. However, as I will detail below, these observations are based on misleading or incomplete reporting of data.

Firstly, the authors compared pairwise topological differences between the trees inferred under six different model selection criteria and reported 0–26% incongruently inferred topologies, depending on the criteria assessed and the dataset employed (their Fig. 1). While it is debatable if this level of incongruence constitutes a “marginal impact on the resulting tree topology”^1^, the most striking trend from these comparisons was not addressed: Across all datasets, differences in topologies between any two best models are considerably lower than distances between a fixed model (GTR+I+G4 or JC) and a best model (Fig 1.). Consistently, all model selection criteria result in very similar trees, which however are fairly dissimilar to trees reconstructed without prior model selection. While these comparisons do not take “accuracy” into account, they are compatible with previous studies finding that any form of model selection results in more accurate topologies compared with using a fixed model^2,3^.

**Fig. 1.**
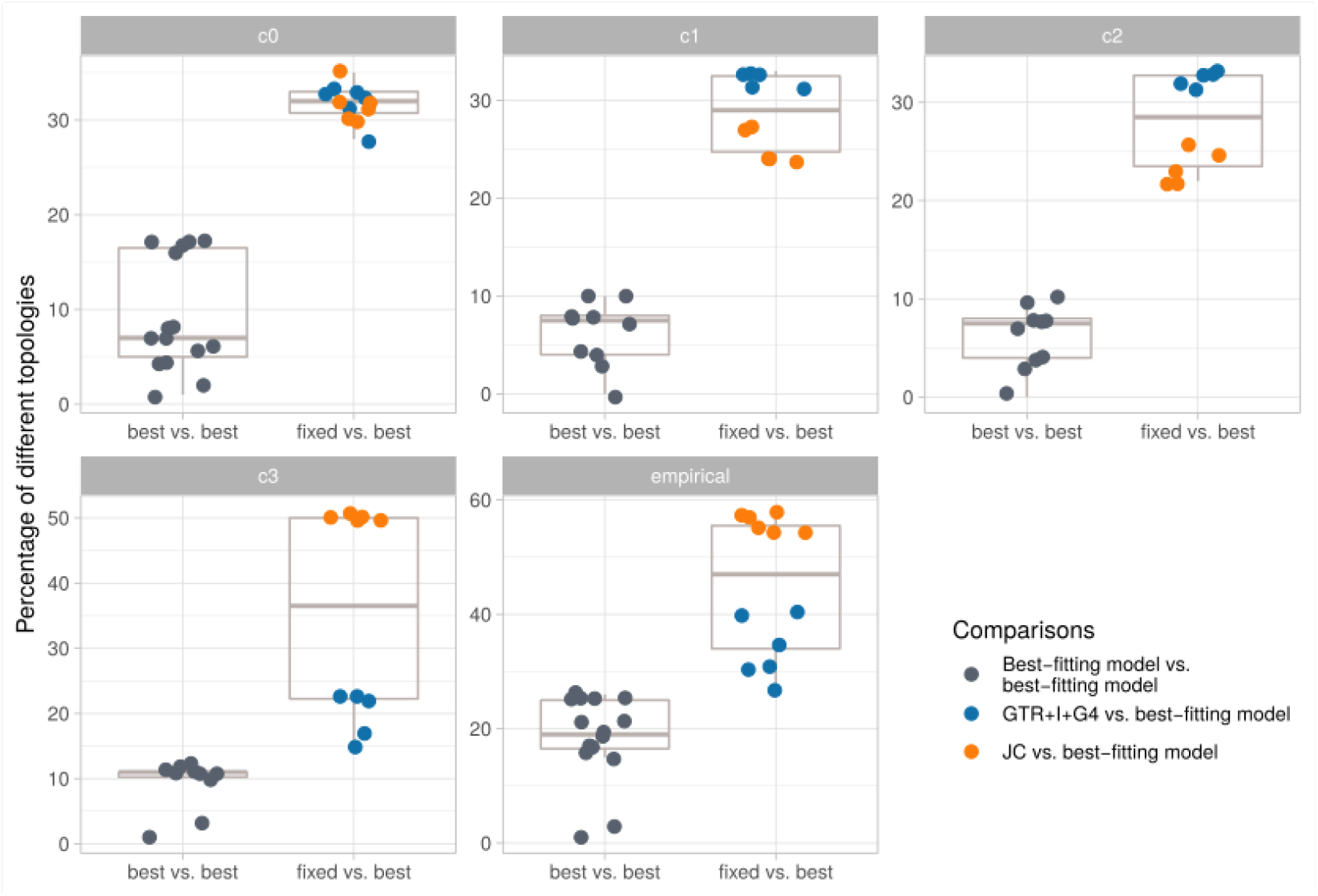
Pairwise comparisons between topologies inferred after model selection or a fixed model. The percentage of differently inferred topologies is plotted, grouped by comparisons between best models inferred by a model selection criterion and comparisons between a best model and a fixed model. Each plotted point represents one comparison, and the panels correspond to the different datasets. All data taken from Fig. 1 in Abadi et al.^1^.

Secondly, the authors counted the number of trees inferred with best, fixed, and true models that are identical to the “true tree”, and found that on average ~50% of trees are correctly inferred by any model or criterion (their Table 2). This representation is problematic, as it does not account for differences in incorrectly inferred trees, and more importantly, averages over all true models. While on average the proportion of correctly inferred trees may be similar, it is unclear if the similarities are consistent across all 7,200 alignments, or if certain selection criteria perform better or worse under certain alignment properties. To address this issue, I have re-analysed the empirical dataset. Maximum likelihood tree reconstructions were performed for all alignments under GTR+I+G4 and under a best model determined using BIC. Both approaches resulted in identical topologies for ~60% of the alignments, which is in agreement with what Abadi et al. found for the empirical dataset (their Fig. 1b). However, the proportion of identically inferred topologies strongly depended on the substitution model that best describes the data, and showed a large variation (~30% – >80%, Fig. 2a). Trees from alignments that were best described by simpler models (such as JC and F81) were generally less well recovered by GTR+I+G4 (the most complex of the 24 models investigated), although this trend was not very pronounced (Fig. 2A). This suggests that the characteristics of an alignment are important in determining to what extent GTR+I+G4 can recover the same topology as a best model. Notably, the same can be observed when ignoring differences in nodes that are not statistically supported (Fig. 2b). Although this analysis is based on an empirical dataset, and the true tree is therefore not known, it demonstrates that tree inferences may differ substantially under GTR+I+G4 and an optimal model selected by BIC. This finding agrees with previous studies on empirical and simulated datasets^2,4^.

**Fig. 2.**
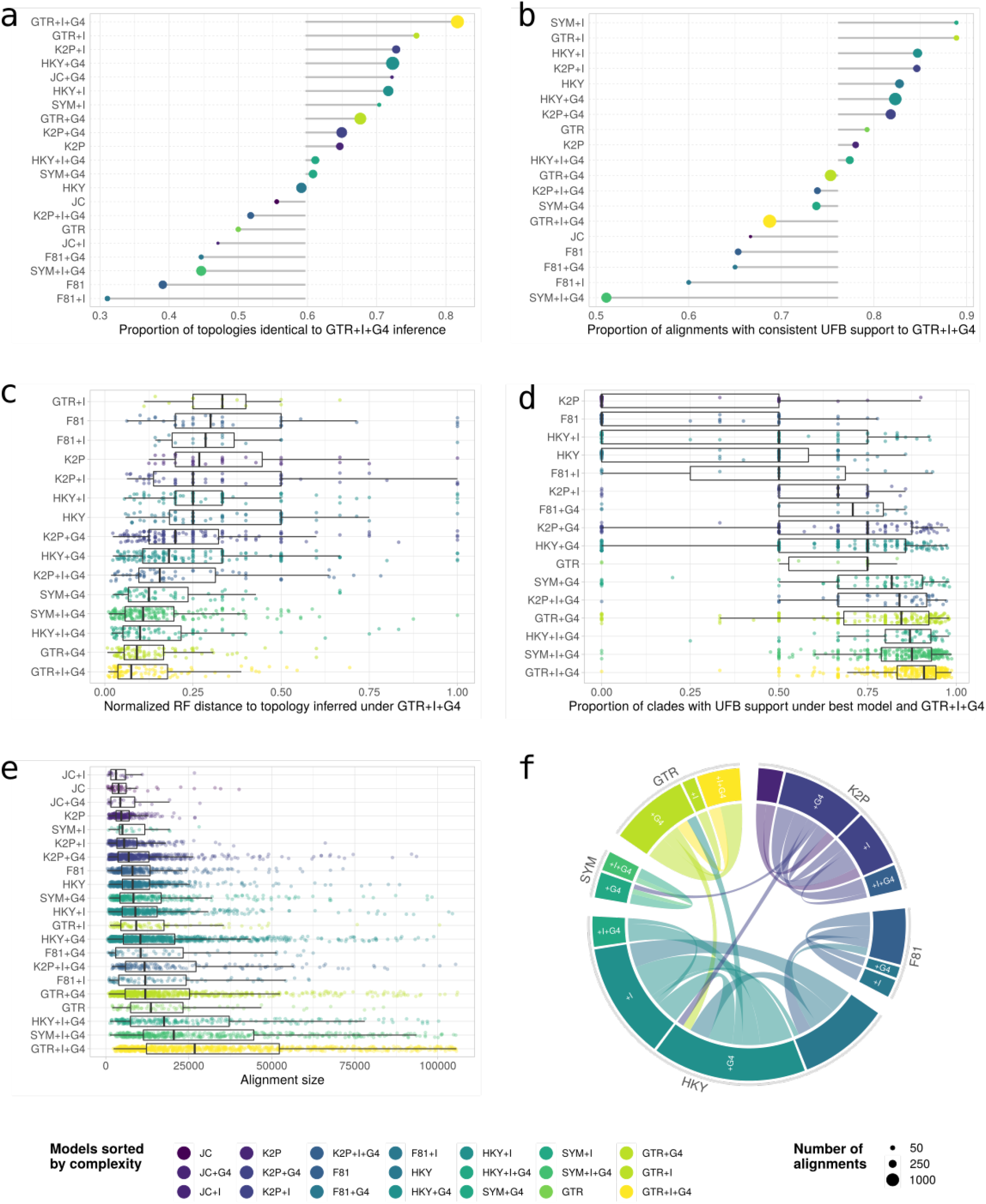
Re-analysis of the empirical dataset. Maximum likelihood trees were reconstructed for all 7200 alignments under a fixed, parameter rich model (GTR+I+G4) and a best model as inferred by BIG. **a** Proportion of identically inferred topologies for each best model compared with GTR+I+G4. **b** Proportion of identically inferred topologies for each best model compared with GTR-I+G4, only considering statistically supported nodes (UFB >= 95), **c** Robinson Foulds distances for non-identical topologies for each best model compared with GTR+1+G4. **d** For non-identical trees, proportion of statistically supported nodes found in trees inferred by a best model and under GTR+I+G4. **e** Alignment size (number of taxa x number of aligned positions) for best models inferred by B1C. **f** Uncertainty in model selection by B1C. Connections in chord diagram represent instances in which multiple models were within the 95% Cl set of the BIC. The size of a connection is relative to how often the two models were within the same Cl set, and the size of sectors is relative to how often each model occurred in any Cl set. To improve visualisation, only connections with at least 100 occurrences in Cl sets are displayed. The total number of displayed connections is 6919.

Thirdly, topological distances between trees obtained under various criteria were reported by the authors to be very similar between all model selection criteria. However, these were either averaged across models and ranked (their Table 3) or binned into 9 categories and averaged (their Fig. 4). Moreover, including distances equal to zero (~50% of all distances) may have obscured patterns in these representations. In my re-analysis, I have therefore investigated topological distances between non-identical trees obtained under GTR+I+G4 and under the best model determined by BIC. Again, distances were inconsistent between alignments, and GTR+I+G4 topologies were most similar to topologies obtained under more complex best models (Fig. 2c). This pattern can also be observed when considering only statistically supported nodes (Fig. 2d).

In summary, the authors’ own data and the here presented re-analysis comparing the best model under BIC with GTR+I+G4 provide compelling evidence that model selection does affect phylogenetic inference. While using GTR+I+G4 produces identical or very similar topologies to any best model identified by a model selection criterion in most cases, the degree of similarity strongly depends on the properties of the underlying alignment: for those alignments that are best described by simple, parameter-poor evolutionary models, GTR+I+G4 often produces very different, but statistically supported phylogenetic estimates (Fig 2a-d). For the empirical dataset, the complexity of the best model chosen by BIC seemed to positively correlate with the size of the dataset (Fig 2e). This suggests that consistently using a fixed parameter-rich model is especially inappropriate for smaller alignments (few taxa and/or few aligned positions).

Overall, the findings discussed here are in agreement with what seems to be a consensus of the literature: There are nuanced differences between model selection criteria^5–7^, but model selection is generally beneficial for phylogenetic accuracy^8–10^.

In addition to inappropriate averaging over alignments with divergent properties, other factors might explain why Abadi et al. did not find differences between the investigated model selection criteria. For example, although a single best model is selected by each of the criteria, other models often cannot be rejected with confidence. In the empirical dataset, the 95% confidence set of BIC supported more than one model for ~79% (5695/7200) of the alignments (Fig. 1f). Taking into account overlapping confidence intervals of different model selection criteria might reduce spurious differences in model choice between the criteria potentially observed by Abadi et al.․ Another factor that should be accounted for in future investigations is tree shape. Ripplinger and Sullivan^11^ found that model fitting is more important when tree stemminess is low. In line with this, for the empirical dataset, topological distances between GTR+I+G4 and the best model inferred by BIC correlated with the proportion of small internal nodes (here defined as internal nodes shorter than 0.1% of the tree length, R^2^=0.6, p < 2.2e-16).

In conclusion, while GTR+I+G4 very often results in accurate phylogenetic estimates even when it is not the best fitting model, its performance is inconsistent across empirically determined alignment properties. There is a large body of literature illustrating the benefits of model selection to phylogenetic inference (reviewed in reference 10). The data presented by Abadi et al. do not provide a convincing justification for skipping model selection. Since convenient and accurate approaches to model selection for maximum likelihood phylogenetics exist^12,13^, the current practice of model selection is not computationally prohibitive. Importantly, only a very limited number of nucleotide substitution models was discussed here. As the field of phylogenetics moves towards larger datasets and increasingly realistic models^14,15^, model selection and fitting will likely become more relevant in the future.

## Methods

The empirical alignments were obtained from https://doi.org/10.17605/OSF.IO/T3PF2. All maximum likelihood analyses were done with IQ-TREE version 1.4.2.^16^, and support estimated with 1,000 ultrafast bootstrap replicates^17^. Best models were determined by BIC under full tree searches for all models and alignments with ModelFinder^13^ implemented in IQ-TREE.

## Acknowledgements

This work would not have been possible without funds from the Johnston Researcher Development Fund of the Institute of Integrative Biology of the University of Liverpool.

## Author contributions

MG conceived the work, analysed and interpreted data, and wrote the manuscript.

